# NMR solution structure and analysis of isolated S3b-S4a motif of repeat IV of the human cardiac sodium channel

**DOI:** 10.1101/2021.05.21.443843

**Authors:** Adel K. Hussein, Mohammed H. Bhuiyan, Boris Arshava, Jianqin Zhuang, Sébastien F. Poget

## Abstract

Voltage-gated sodium channels are membrane proteins that play an important role in the propagation of electrical signals by mediating the rising phase of an action potential. Numerous diseases, including epilepsy, extreme pain, and certain cardiac arrhythmias have been linked to defects in these channels. The S3b-S4a helix-turn-helix motif (paddle motif) is a region of the channel that is involved in voltage sensing and undergoes significant structural changes during gating. It is also the binding site for many gating-modifier toxins. We determined the solution structure of the paddle motif from the fourth repeat of NaV1.5 in dodecylphosphocholine micelles by NMR spectroscopy and investigated its dynamics and micelle interactions. The structure displays a helix hairpin with a short connecting loop, and likely represents the activated conformation with three of the first four gating charges facing away from S3. Furthermore, paramagnetic relaxation measurements show that the paddle motif is mainly interacting with the interface region of the micelle. NMR relaxation studies reveal that the paddle motif is mostly rigid, with some residues around the loop region and the last 4 residues on the C-terminus displaying heightened mobility. The structural findings reported here allowed the interpretation of three disease-causing mutations in this region of the human cardiac sodium channel, S1609W, F1617del and T1620M. The establishment of this model system for NMR studies of the paddle region offers a promising platform for future toxin interaction studies in the cardiac sodium channels, and similar approaches may be applied to other sodium channel isoforms.

---

Voltage gated sodium channels (VGSCs) are membrane proteins that function in the propagation of electrical signals by depolarizing excitable cells^1,2^. Numerous diseases have been linked to mutations in VGSCs including epilepsy, mental retardation, and several ailments that could lead to sudden cardiac death^3^. There are nine well characterized sodium channel isoforms, NaV1.1-1.9, that are expressed in a tissue specific manner. NaV1.5 is expressed mainly in cardiomyocytes by the gene SCN5A and is involved in the electrical signaling of the heart^4^. Many diseases are associated with mutations in SCN5A, such as Brugada syndrome, long-QT syndrome and sick sinus syndrome^5^. While the heart expresses other sodium channel subtypes, they cannot compensate for the dysfunction in NaV1.5, which demonstrates the importance of NaV1.5 to the function of cardiac cells. Furthermore, NaV1.5 is a major target of class I antiarrhythmic drugs, such as lidocaine^3^.

Mammalian VGSCs consist of a pore forming α subunit and auxiliary β subunits. The α subunit consists of one polypeptide chain with 24 transmembrane segments organized into four repeats (domains) that surround a central aqueous pore^2,3,6^. Each repeat consists of six transmembrane helical segments labeled S1-S6. The S1-S4 segments make up the voltage sensing domain (VSD), while the S5-S6 and the extracellular loops (p-loops) that connect these two segments from all 4 repeats collectively constitute the pore domain^2^. Much information on the structure of VGSCs has been obtained from the structure elucidation of bacterial sodium channels such as Na_V_AP^7^, Na_V_RH^8^, Na_V_Ct^9^ and Na_V_Ms^10^, and more recently from the cryo-EM structures of the Na_V_1.2^11^, Na_V_1.4^12^, Na_V_1.5^13^, and Na_V_1.7^14^ mammalian channels. The S4 helix is known as the voltage sensor and contains basic residues (typically arginines) at every third position^2^. During resting potential, the S4 voltage sensor is facing downward towards the intracellular side, and during depolarization it moves outward towards the extra-cellular side^2^. Movement of the S4 voltage sensor in repeats 1-3 translates to channel activation, and in repeat 4 to channel inactivation^2,15^.

Voltage gated sodium channels are the targets of certain animal toxins. These toxins are an excellent probe of channel function and a good drug lead due to their specific and potent effect. Toxins that target the VSD usually bind to a region of the S3 and S4 helices known as the S3b-S4a or “paddle” motif^3^. The paddle motif is a modular unit that is also implicated in the gating process^16^. It was demonstrated that the paddle motif can be transferred between different channels and retain its function and toxin binding^17,18^. The kinetics of VSD4 have been shown to be slower than the other repeats^19^, indicating that VSD4 plays a role in determining the rate of channel inactivation^15,18^. Consequently, many toxins that interact most strongly with domain IV paddle motif were shown to slow down inactivation^18^. Considering that certain heart diseases are associated with accelerated fast inactivation, NaV1.5 domain IV paddle motif represents a good drug target^20,21^.

Although the structure of Na_V_1.5 was recently determined by cryo-electron microscopy^13^, the resolution in the region of the paddle motif is between 4 and 6 Å. Furthermore, the detergent that was used for the reconstitution of the full channel does not have a phosphate head group. This could lead to some structural distortions since it is known that interactions between the VSDs and the lipid environment are important for proper function and toxin binding^16^.

In this study, a simple and efficient method was devised to obtain high yields of isotopically labeled paddle motif from NaV1.5 DIV. Furthermore, we determined the solution NMR structure of NaV1.5 DIV paddle motif in DPC micelles. This paddle motif consists of 37 residues with a sequence of VVVILSIVGTVLSDIIQKYFFSPTLFRVIRLARIGRI, which corresponds to residues 1597-1633 of human Na_V_1.5. Using paramagnetic relaxation enhancement, we investigated the positioning of each residue with respect to the micelle. Backbone dynamics of the paddle motif in DPC micelle were determined through NMR ^15^N relaxation measurements.

## Materials and methods

### Materials

n-Decyl- and n-dodecylphosphocholine (Fos-Choline-10 and DPC) were obtained from Anatrace Products (Maumee, OH). cis-[Pt(en)Cl_2_] (en = ethylenediamine) was obtained from Alfa Aesar (Haverhill, MA). The lipids 1,2-dimyristoyl-sn-glycero-3-phosphocholine (DMPC) and 1,2-dihexanoyl-sn-glycero-3-phosphocholine (DHPC) were obtained from Avanti Polar Lipids (Alabaster, AL).

### Cloning, expression, and inclusion bodies isolation of Thioredoxin-paddle fusion protein

A codon-optimized gene fragment coding for residues 1597-1633 of NaV1.5 preceded by a methionine residue and followed by a stop codon was cloned into pET-32b(+) (Novagen) between the BglII and XhoI restriction sites using standard molecular biology techniques. The resulting plasmid encodes a thioredoxin-paddle fusion protein separated by a 47 amino acid linker containing a hexa-histidine tag (Figure S1). Fusion protein was over-expressed in *E. coli* BL21(DE3) C43 cells^22^ at 37 °C. Cells were induced at an optical density of 0.4 with 1 mM isopropyl β-d-1-thiogalacto-pyranoside (IPTG) and grown for 18 hours post induction at 37 °C. Inclusion bodies were isolated based on the method by Cohen *et al.*^23^ with minor modification. Briefly, cells were spun down at 3220 × g for 20 minutes and cell pellets were lysed by suspending in lysis buffer (50 mM Tris pH 8.0, 0.5 mM EDTA) containing 10 mg/mL lysozyme followed by sonication. Inclusion bodies were isolated by spinning the cell lysates at 38,700 × g at 4 °C. The resulting pellet was washed three times by resuspension followed by centrifugation as before, first with 50 mM Tris buffer pH 8.0, then Tris buffer supplemented with 2 % triton-X 100, and finally pure water.

### Purification of paddle peptide

The cleavage reagent complex cis-[Pt(en)(H_2_O)_2_]^2+^ was generated by reacting cis-[Pt(en)Cl_2_] with silver nitrate in a 1:2 molar ratio in water (pH 4.0) at 42 °C for 2 h. Inclusion bodies containing thioredoxin-paddle fusion protein were dissolved in 80 % formic acid and cleaved by the addition of 5 mM of this complex and incubation at 60 °C for 6 hours. The cleaved peptides were precipitated by the addition of sodium hydroxide until the pH reached a value of 8.0. The precipitated mixture was dissolved in 6 M guanidium chloride, 0.5 M HCl, incubated for 16 h at 4 °C and purified by HPLC on a Zorbax C3 column with dimensions of 21 × 150 mm (Agilent, Santa Clara, CA) using a gradient of 20-90 % acetonitrile in water with 0.1 % trifluoroacetic acid (TFA) in 60 min at a flow rate of 5 ml/min. Eluted peptide was lyophilized, dissolved in 1:1 trifluoroethanol (TFE):water, lyophilized again and stored at −20 °C.

### NMR sample preparation

Lyophilized paddle motif was dissolved in buffer containing 10 mM potassium phosphate, pH 7.0, 10 mM NaCl, 0.05 % sodium azide, 10 % D_2_O, 50 mM Fos-Choline-10 or 100 mM DPC. For bicelle, lyophilized paddle motif, DMPC and DHPC were dissolved in 1:1 chloroform:methanol, dried under nitrogen gas and dissolved in buffer containing 10 mM potassium phosphate, pH 7.0, 10 mM NaCl, 0.05 % sodium azide, 10 % D_2_O. Total lipid concentration was 300 mM with *q*=0.5 (*q*=long chain:short chain molar ratio). All spectra were recorded at 37 °C on either Agilent (Varian) NMR system 600 MHz spectrometer (paramagnetic, and homonuclear NOESY, TOCSY experiments) or 800 MHz Bruker Avance (all other experiments), both equipped with a triple resonance cold probe. All NMR spectra were processed using NMRPipe^24^ software and analyzed on CCPNMR^25^ analysis software.

### Backbone assignment in Fos-Choline 10

Chemical shifts were referenced to sodium 2,2-dimethyl-2-silapentane-5-sulfonate (DSS), and ^15^N and ^13^C shifts were referenced indirectly^26^. For backbone assignments, HNCA, HNCACB, HNCOCA and HNCO triple resonance experiments were collected on a ^13^C^15^N uniformly labeled peptide at a concentration of 500 μM. The parameters are as follows: HNCA (time domain data size 64(*t1*) × 40(*t2*) × 1024(*t3*) complex points; spectral width 30, 26, and 15.9 ppm for (*t1*), (*t2*) and (*t3*) respectively; 40 scans), HNCACB (time domain data size 128(*t1*) × 40(*t2*) × 1,024(*t3*) complex points; spectral width 70, 26, and 15.9 ppm for (*t1*), (*t2*) and (*t3*) respectively; 64 scans), HNCO (time domain data size 56(*t1*) × 40(*t2*) × 2,048(*t3*) complex points; spectral width 14, 26, and 13.9 ppm for (*t1*), (*t2*) and (*t3*) respectively; 8 scans), HNCOCA (time domain data size 64(*t1*) × 40(*t2*) × 2,048(*t3*) complex points; spectral width 30, 26, and 16 ppm for (*t1*), (*t2*) and (*t3*) respectively; 24 scans).

### Backbone assignment in DPC

Assignments were obtained through analysis of HNCA, HNCACB and ^15^N NOESY experiments collected on ^13^C^15^N uniformly labeled peptide at a concentration of 500 μM. Side chains were assigned using a combination of ^15^N NOESY, ^13^C NOESY, and HCCH-TOCSY experiments. The following experimental parameters were used: HNCA (time domain data size 64(*t1*) × 50(*t2*) × 1024(*t3*) complex points; spectral width 32, 20, and 12 ppm for (*t1*), (*t2*) and (*t3*) respectively; 32 scans), HNCACB (time domain data size 120(*t1*) × 50(*t2*) × 1,024(*t3*) complex points; 16 scans), ^15^N NOESY (time domain data size 128(*t1*) × 48(*t2*) × 1,024(*t3*) complex points; 16 scans, mixing time = 150 ms), ^13^C NOESY (time domain data size 200(*t1*) × 50(*t2*) × 1,024(*t3*) complex points;16 scans, mixing time = 150 ms), HCCH-TOCSY (time domain data size 180(*t1*) × 80(*t2*) × 1,536(*t3*) complex points; 16 scans, spin-lock period = 100 ms).

### Structure calculation

Dihedral angles were obtained from TALOS-N^27^ using the Cα, Cβ, C, and Hα chemical shifts. Distance restraints were obtained from the ^15^N NOESY and ^13^C NOESY experiments. All NOESY peaks were assigned manually. Structure calculation was done through ARIA2.3^28^/CNS1.21^29^ standard protocol using the NOE distance and the TALOS dihedral angle restraints. Distances were calibrated by cross peak volume within the software. The structural calculations consisted of 8 iterations of simulated annealing. The first 6 iterations generated 20 structures each, while the last 2 iterations generated 100 structures each with the 7 lowest energy structures chosen as input for the next iteration at each step. The 10 lowest energy structures of the last iteration were water refined. Structure quality was assessed by PROCHECK-NMR^30^. All structure calculations were carried out using the NMRBox virtual software environment^31^.

### Paramagnetic relaxation

Paddle motif at a concentration of 300 μM was prepared with and without 5-doxyl stearate (Sigma-Aldrich, St. Louis, MO) or 16-doxyl stearate (SigmaAldrich) in the same DPC buffer as above at 1:2 peptide:doxyl stearate molar ratio. For the manganese experiment, paddle motif and MnCl2 were dissolved in 10 mM HEPES, pH 7.0, 10 mM NaCl, 100 mM DPC. Samples were prepared at 1 mM paddle concentration, and HSQC spectra were measured before and after addition of 0.5 mM Mn^2+^. Peak height was normalized to the least affected residue.

### NMR relaxation

Relaxation parameters (*R*_1_, *R*_2_ and heteronuclear NOE) were measured on an 800 MHz spectrometer using the standard ^15^N HSQC based pseudo 3D experiments on a 1 mM ^15^N-labeled paddle sample dissolved in 100 mM DPC buffer. For *R*_1_, nine interleaved experiments with the following relaxation delays were collected: 0, 160, 240, 400, 560, 720, 880, 960, and 1040 ms. For *R*_2_, six interleaved experiments were collected with relaxation delays: 0, 16, 32, 64, 96, and 128 ms. HetNOE were collected using a saturation time of 4 s for the saturation experiments, and an overall relaxation delay of 7 s. Four HetNOE experiments were measured in interleaved fashion, and the average values were obtained from two experiments (two for saturation, and two for non-saturation) with the uncertainty determined as the average standard error. All analysis was done on CCPNMR analysis software. The S^2^ order parameters were obtained from Lipari-Szabo model free analysis^32^ using the FAST-Modelfree program^33^.

## Results and discussion

### Sample preparation

For initial experiments, paddle peptide was expressed in inclusion bodies as TrpΔLE fusion and the fusion tag cleaved off with cyanogen bromide due to the success of other labs in expressing membrane protein fragments using this system^23^. However, due to the health hazards of working with cyanogen bromide, low yield, and occasional occurrences of peptide oxidation, we developed a safer and more reliable peptide production protocol involving expression as a thioredoxin fusion protein and cleavage by *cis*-[Pt(en)(H_2_O)_2_]^2+^ complex. This complex cleaves at the C-terminus of methionine residues just like cyanogen bromide^34^. Optimization of cleavage conditions showed essentially complete cleavage of fusion protein after a 7 hour incubation with 5 mM platinum complex at 60 °C (Figure 1). Analysis of the cleavage reaction by MALDI-TOF mass spectrometry confirmed the presence of the cleaved paddle peptide. In addition to the paddle motif, the mass spectrum also shows two more peaks that correspond to formylated peptides. Formyl groups were removed by incubation with 0.5 M HCl before HPLC purification (Figure 2). The cleavage mixture was purified by preparative HPLC and purity was assessed by analytical HPLC (Figure S2). This method resulted in an improved peptide yield of 10-15 mg of peptide per liter of growth, compared to the 2-4 mg yield achieved by cyanogen bromide cleavage of TrpΔLE fusion construct, and also avoids the presence of residues modified by oxidation in the final peptide product.

**Figure 1:**
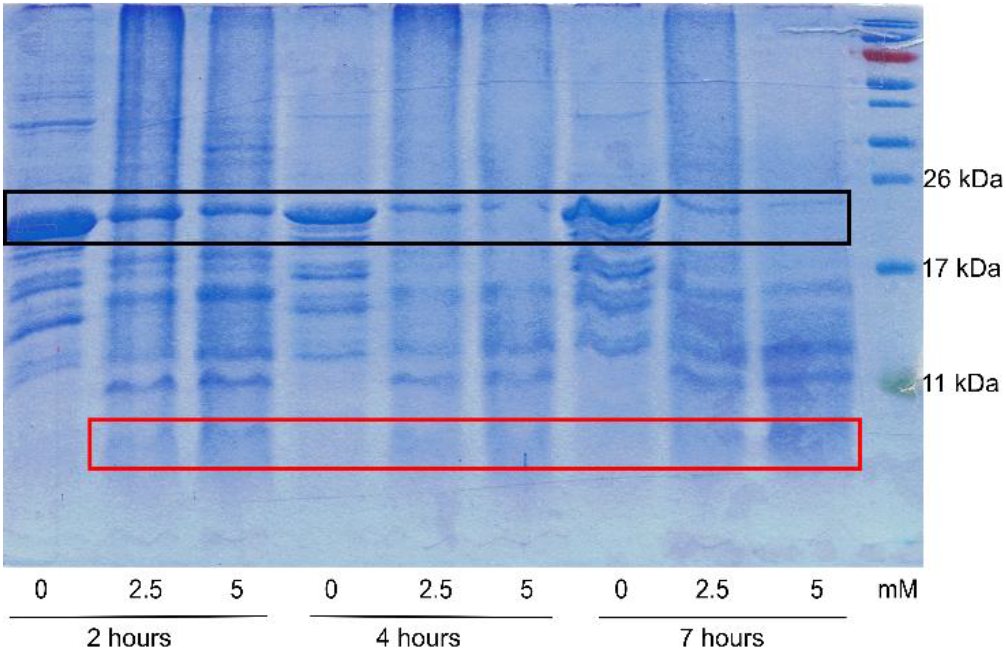
SDS-PAGE analysis of the fusion protein cleavage by *cis*-[Pt(en)(H2O)_2_]^2+^ complex. Redissolved inclusion bodies were cleaved with varying platinum complex concentrations for different amounts of time, and reaction completion analyzed on a 16 % TRIS-Glycine gel. Note the disappearance of the fusion protein (the bands just below the 26 kDa marker region in the black box) and appearance of low molecular weight peptide (in the red box) following incubation with the platinum complex.

**Figure 2:**
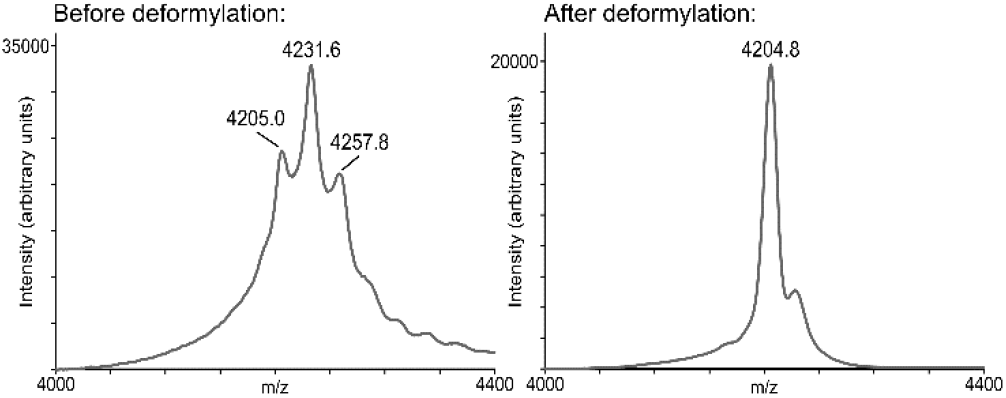
Deformylation of paddle motif following cleavage in 80 % formic acid with *cis*-[Pt(en)(H2O)_2_]^2+^. (left) A MALDI-TOF mass spectrum shows three major peaks representing unformylated, singly formylated and doubly formylated paddle motif. The expected mass of the paddle motif is 4205 Da. The two peaks with about 27 Da and 54 Da higher mass correspond to the presence of one or two formyl group adducts. (right) Paddle motif after reaction with 0.5 M HCl for 16 h at 4 °C and purification on HPLC shows the disappearance of most of the formylated adducts and a single peak at 4204.8 Da that matches the theoretical paddle mass.

While the expression of transmembrane fragments as TrpΔLE fusion proteins followed by cyanogen bromide cleavage to liberate the transmembrane region is a commonly used procedure^35^, we found that expression as a thioredoxin fusion protein with platinum complex cleavage had several advantages. Foremost, the overall safety of the procedure was increased by the replacement of the cyanogen bromide cleavage step. Additionally, the thioredoxin fusion construct resulted in superior expression levels as well as purity of the inclusion bodies. We found that using the standard method, the TrpΔLE fusion protein needed to be washed several times and purified using Ni^2+^ affinity resin to obtain sufficient purity for chemical cleavage, whereas the redissolved thioredoxin inclusion bodies could be cleaved directly without prior purification, simplifying the overall protocol. This method therefore may provide an alternative route for producing high quantities of transmembrane protein segments for NMR and other biochemical studies.

### NMR assignments

Backbone assignments of the peptide in 50 mM Fos-Choline-10 were made by collecting HNCA, HNCACB, HNCOCA, and HNCO triple resonance experiments on a ^13^C^15^N double labeled sample and sequentially linking chemical shifts starting from glycines at positions 1605 and 1631 (Figure S3). Full assignments were obtained for all backbone hydrogen, nitrogen, and carbon resonances with the exceptions of the first two N-terminal amino acids, which could not be observed in any spectra, and for proline 1619 and threonine 1620, for which only the Cα atoms were assigned.

HSQC spectra of paddle motif in 100 mM DPC were highly similar to those in Fos-Choline-10 (Figure S4), and therefore about half of the assignments could be unambiguously transferred to the DPC spectra. Using those peaks as starting points, ^15^N NOESY, HNCA, and HNCACB spectra were used to assign the rest of the peaks. Assignments in DPC were obtained to the same extent as in Fos-Choline-10 (Figure 3). In addition to these assignments in detergent, a sample of paddle peptide in DHPC/DMPC bicelles (with a 2:1 molar ratio of short-chain to long-chain lipid) was also prepared, and the resulting HSQC spectrum showed only minor differences in peak position to the spectra in detergent (Figure S5), indicating that the conformation in detergent micelles is very close to that in the more membrane-like bicelle environment.

**Figure 3:**
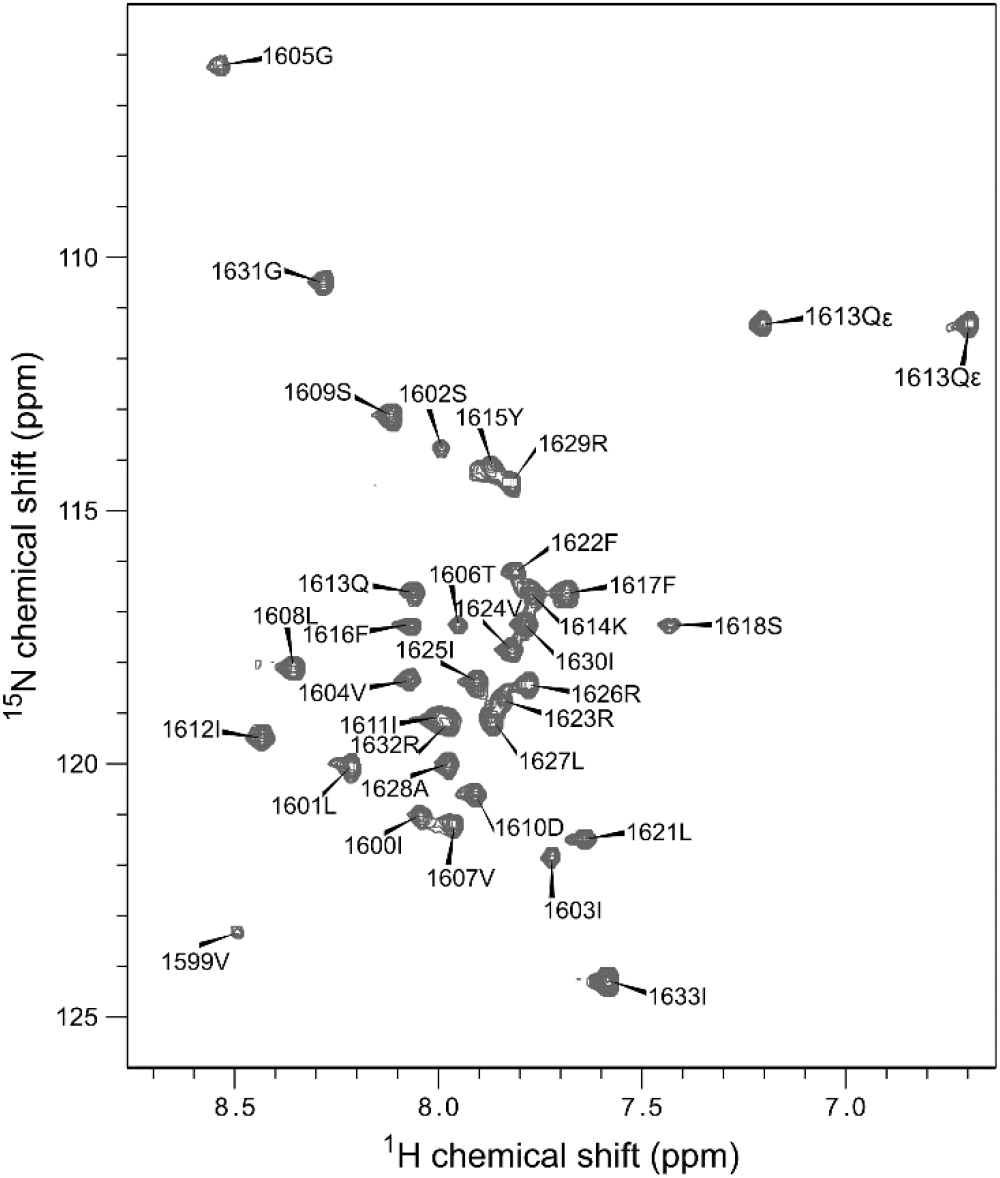
Fingerprint of NaV1.5 DIV paddle motif. Assigned ^15^N HSQC spectrum of paddle motif in DPC micelles.

Side chain assignments in DPC micelles were based on ^15^N NOESY, ^13^C HCCH-TOCSY, ^13^C NOESY (in D_2_O and H_2_O) and HNCACB spectra collected on a sample prepared with per-deuterated detergent. Hα chemical shifts were identified based on the backbone assignments with help of the ^15^N NOESY spectrum, and the rest of the side chain proton and carbon assignments relied on the HCCH-TOCSY and HNCACB experiments. About 85% of the side chain protons were assigned using this approach. Unassigned side chains include the first two residues and Leu 1608. For the latter, side chain resonances could not be assigned due to overlap in spite of existing backbone assignments.

### Structure of S3b-S4a paddle motif in DPC micelles

The structure was calculated using dihedral angles obtained from the chemical shifts and distance restraints from the ^15^N NOESY and ^13^C NOESY spectra. The first structure calculation was performed using the dihedral angle restrains and unambiguous distance restraints. The resulting structure ensemble was used to assign some of the ambiguous distance restraints and a second round of structural calculation was performed. Initial structure calculations established the helix-coil-helix motif. After several rounds of structural refinement, the final 10 lowest energy, water refined structures converged (Figure 4, see Table 1 for structural statistics). Several long-range NOEs proved crucial for determining the overall structure. They include an NOE between 1612IleHα to 1616PheHδ and 1612IleHδ to 1622PheHδ.

**Figure 4:**
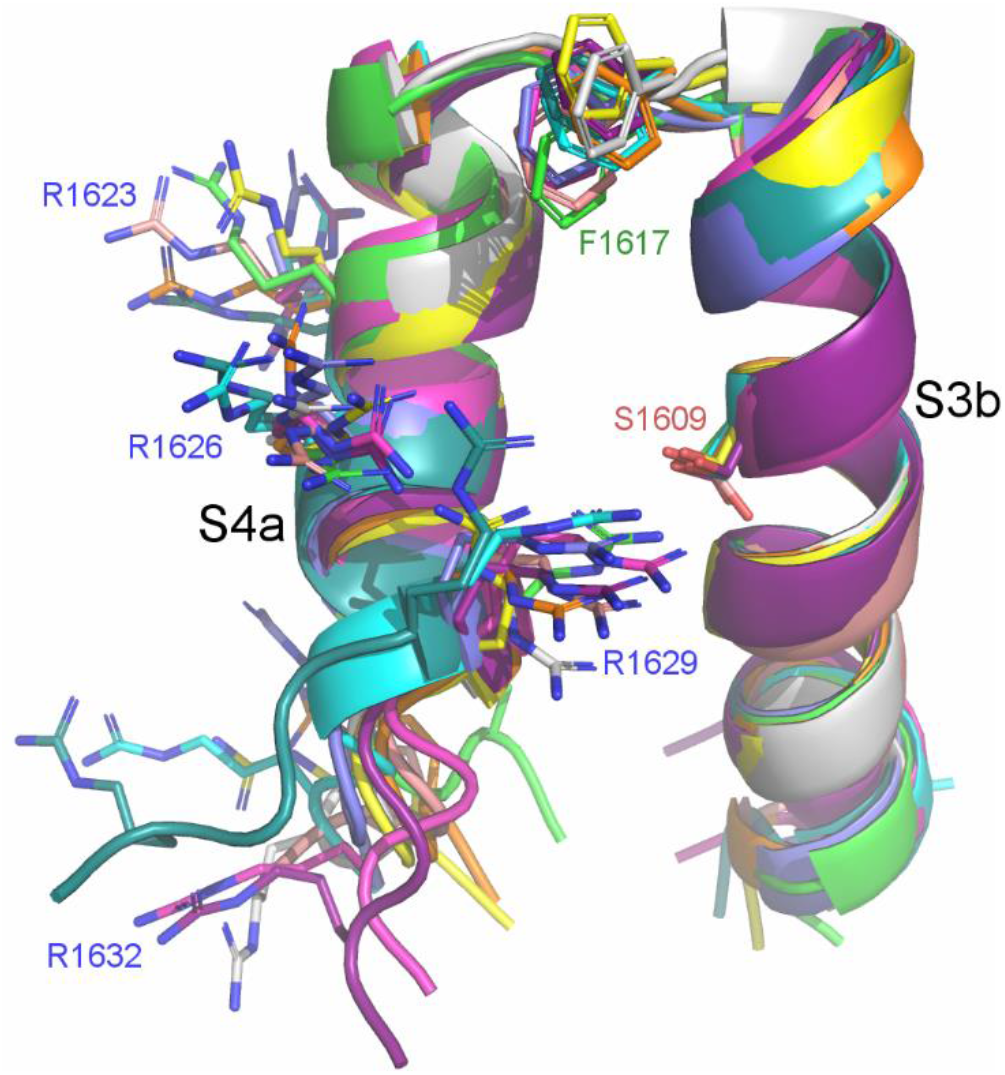
Overlay of the 10 lowest energy water refined structures of NaV1.5 DIV S3b-S4a paddle motif generated by Aria2 shown as a cartoon representation. Functionally important residues discussed in this manuscript are shown in stick representation (hydrogen atoms omitted for clarity).

**Table 1:** Structural statistics for the final 10 models of NaV1.5 DIV S3b-S4a.

| Quantity | Value |
| --- | --- |
| Total unambiguous distance restraints | 755 |
| Intra residue | 606 |
| Sequential ($\vert i - j\vert = 1$) | 68 |
| Medium ($2 \leq \vert i - j\vert \leq 4$) | 73 |
| Long range | 8 |
| NOE violations (*mean and s.d.*) |  |
| $> 0.5 \text{ \AA}$ | 5.8±0.6 |
| $> 0.3 \text{ \AA}$ | 12.9±1.51 |
| $> 0.1 \text{ \AA}$ | 44.4±2.90 |
| RMSD from the average atomic coordinates ($\text{\AA}$) | |
| Backbone |  |
| 2<sup>nd</sup> structure | 0.559±0.131 |
| All residues | 1.47±0.521 |
| Heavy atoms |  |
| 2<sup>nd</sup> structure | 1.12±0.119] |
| All residues | 1.88±0.450 |
| Deviations from idealized covalent geometry |  |
| Bond ($\text{\AA}$) | 0.00507±0.000166 |
| Angles ($^\circ$) | 0.59 ± 0.024 |
| Improper dihedrals ($^\circ$) | 1.73 ± 0.11 |
| Ramachandran analysis (%) |  |
| Residues in most favored regions | 81.2 |
| Residues in additionally allowed regions | 18.8 |
| Residues in generously allowed regions | 0.0 |
| Residues in disallowed regions | 0.0 |

As expected, the structure consists of mostly alpha helix. The only exceptions are the termini and the loop. The lack of order in the N-terminal region is due to the lack of assignments of the first two residues. Even though all residues in the C-terminus were assigned, there was a lack of inter-residue NOEs in this region. This could be due to increased flexibility in this region, which is supported by NMR relaxation studies (Figure 8). A certain level of unraveling of the alpha helices at both termini would be expected due to the fact that the analyzed peptide is a truncated fragment missing the intracellular halves of the S3 and S4 helices. However, the significant difference in the extent of that unraveling at the N- and C-terminal ends also indicates sequence-specific differences that may have functional relevance (further discussed below). When comparing the conformation of the side chains for arginine residues 1623, 1626, 1629 and 1632 (the gating charge residues R1 to R4), it can be seen that R1 to R3 side chain positions are quite well defined in the structural bundle, whereas R4 is disordered due to the unravelling of the S4 helix towards the C-terminus (Figure 4).

While refining the final structure of the paddle motif, the cryo-EM structure of the full rat NaV1.5 channel was published^13^. This provided an opportunity to compare the structure of the paddle motif in isolation and in the context of the full channel. Alignment of the average NMR solution structure of NaV1.5 DIV S3b-S4a with full channel (Figure 5) resulted in a backbone RMSD between the two of 2.9 Å (and of 3.2 Å with the more recently published human NaV1.5 structure^36^), which represents good alignment when compared to the local resolution of ~5 Å in this region of the cryo-EM structure. The high similarity between the solution and the cryo-EM structure is a good validation that the NMR structure of the paddle motif represents a functionally relevant conformation. This is consistent with previous studies that have shown that an isolated paddle peptide retains binding affinity towards toxin even in the absence of the whole channel^37^. Our finding that the Na_V_1.5 paddle motif retains its native structure as an isolated peptide in DPC micelles can explain these previous results.

**Figure 5:**
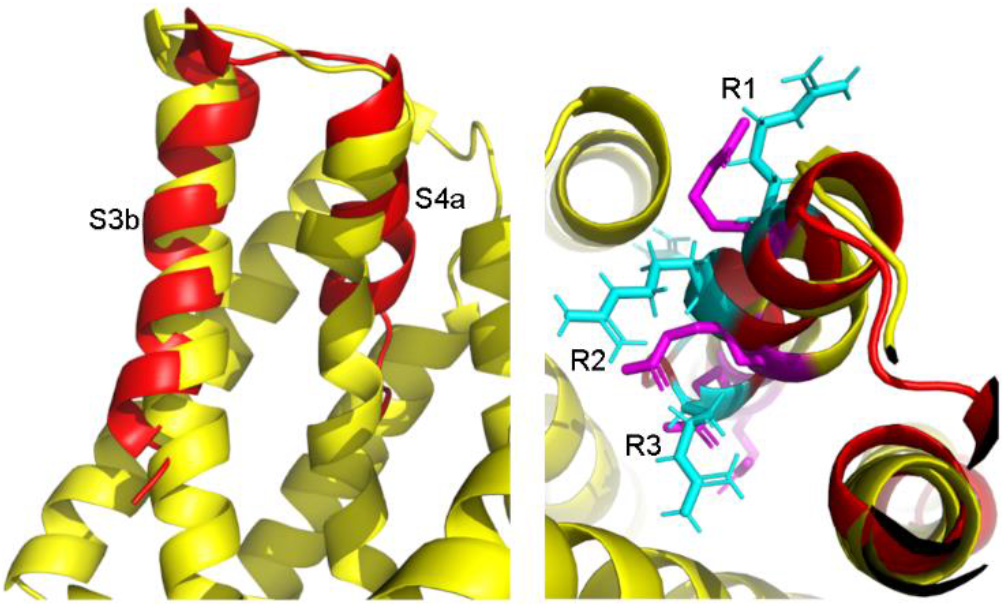
(left) Alignment of NMR solution structure of Na_V_1.5 DIV S3b-S4a motif with cryo-EM of the full channel from rat. (right) Alignment of the gating charges on the S4a helix. Atoms colored in cyan are the arginine residues (gating charges) from the NMR structure while those colored in magenta are from the cryo-EM structure of the full channel.

It is interesting to compare our study with a previous NMR study of a 28 amino acid fragment in DPC micelles representing the S3b-S4 motif of the BK potassium channel. In that study, the potassium channel peptide also adopted a helix-turn-helix motif^38^, although the structure does not align as well with the cryo-EM structure of full BK channel^39^ as our paddle fragment does with full NaV1.5. These differences might be due to the fact that BK channel adopts a much more extensive loop region between the helices, where interactions between different loops that cannot be fully captured in an isolated peptide might contribute to stabilizing the channel structure.

While our paddle structure aligns well with the same region of the full channel structure, a significant difference is found in the S4 helix, which in the cryo-EM structure transitions from an alpha helix to an extended 3_10_ helix after R1. In the NMR structure, the whole S4b region forms a continuous alpha helix up to R3. In the resting state, sodium channel voltage sensors are thought to adopt a 3_10_ helix in the region from R1 to R4, with the residues around R1 relaxing to an α helix upon transition to the activated state^40,41^. In the case of NaV1.7-NavPas chimera cryo-EM structure in the activated state, it was observed that the α-helical conformation extended up to and including the region around R2^42^, but the conformation found in our fragment is unique in the structural studies so far. This could indicate that the 3_10_ helical conformation in S4 is induced to facilitate favorable structural interactions elsewhere in the VSD, and the absence of those interactions allows the peptide to adopt its most relaxed secondary structure seen in our truncated fragment. Alternatively, this difference in secondary structure may also be due to the detergents used in the reconstitution. In our study, the isolated motif was reconstituted in DPC micelle, while in the cryo-EM structure of full Na_V_1.5, the channel was solubilized in glycol diosgenin (GDN) supplemented with cholesteryl hemisuccinate. The presence of choline phosphate headgroups in DPC, which are not present in GDN, may also result in conformational differences. It is known that interaction between ion channels and membrane phospholipids are essential for proper channel function^43^, and some of these interactions involve the paddle motif and influence both toxin binding and gating of the channel^16^. The structure in DPC micelle may therefore be closer to the structure in lipids, since we have observed that the NMR spectra in DPC micelles and DHPC/DMPC bicelles are similar, indicative that the conformation of the paddle in DPC is similar to that in the more membrane-like bicelle system (Figure S5).

### Gating charges

An essential feature of sodium channel VSDs are the gating charges consisting of the positively charged amino acid side chains found at every third position on helix S4. The NaV1.5 DIV paddle motif fragment contains four of these arginine residues that are separated by hydrophobic residues to give it an amphipathic character. In the structure of the paddle motif determined in this report, three of the gating charges (R1, R2 and R4) are facing to the outside of the helical hairpin away from S3, while the third one (R3) points into the hairpin towards the S3 helix (Figure 6). This difference in side chain arrangement is also reflected in the sensitivity to paramagnetic relaxation agents (see below), which show that R3 is the most sensitive to the lipophilic paramagnetic agents, while R1 and R4 are the least sensitive (Figure 7). It should be noted in this context that the region around R4 is unstructured and highly dynamic in the paddle fragment, so our discussion here will focus on R1-R3. During gating, the first two gating charges are facing towards the intracellular side in the resting state and during channel activation, they move outward toward the extracellular side through a “sliding helix” mechanism^13^. In the NMR solution structure of the paddle motif, the first two gating charges are facing outward and the third is facing inward in agreement with other studies. As expected, the NMR solution structure of the S3b-S4a paddle motif resembles the activated state because it was reconstituted in symmetrical DPC micelles that approximate the environment of a membrane at 0 mV potential.

**Figure 6:**
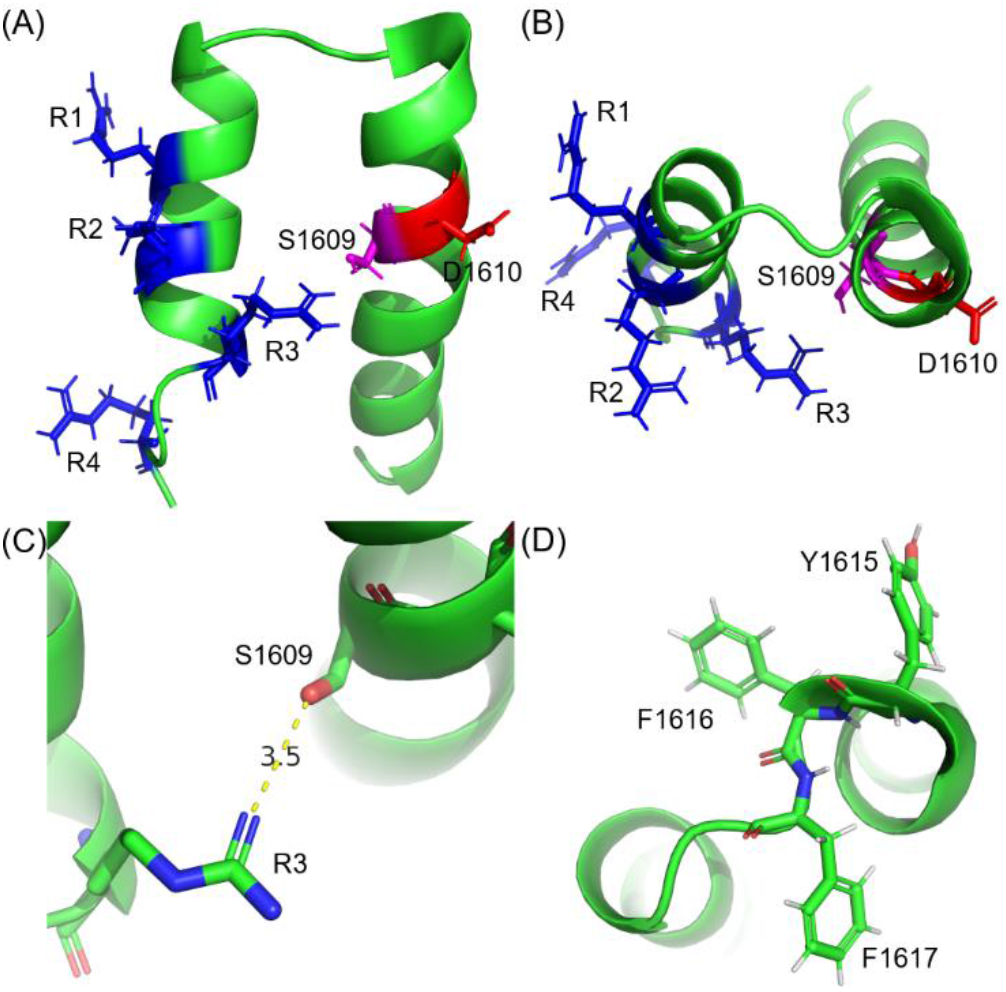
Structural features of NaV1.5 DIV S3b-S4a paddle motif. (A) Top and (B) side views of the paddle motif highlighting the locations and orientation of the gating charges with arginine residues colored in blue, aspartic acid in red and serine in magenta. Three of the gating charges are facing outward, while R3 is facing inward. (C) Detail of the possible interaction between R3 and serine 1609 on the S3b helix. (D) Detail of the loop connecting S3 and S4 helix with aromatic residues shown in stick representation.

**Figure 7:**
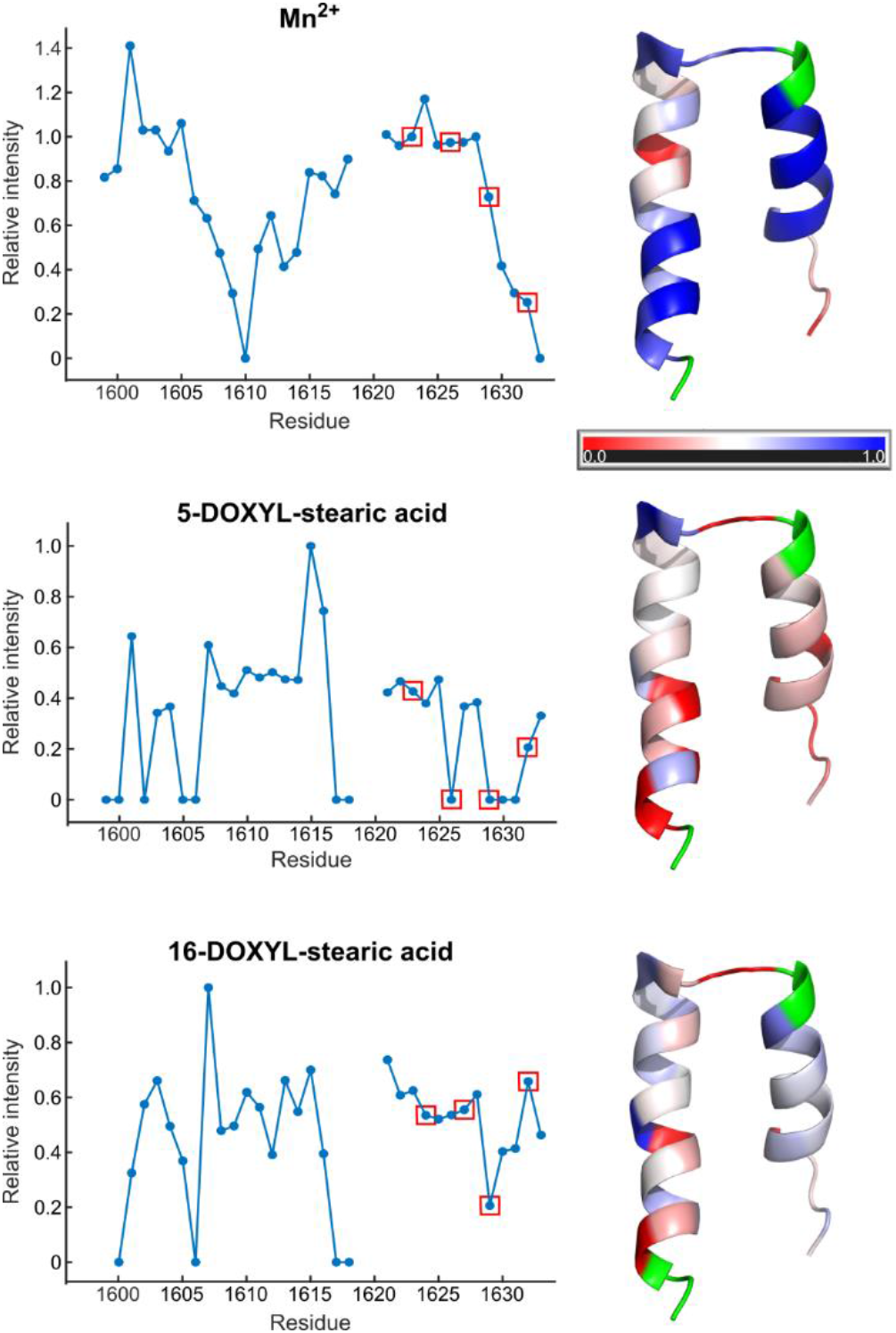
Paddle motif interaction with DPC micelle. The ratios of peak intensity before and after addition of relaxation agent are plotted against residue number for Mn^2+^, 5-DOXYL stearic acid and 16-DOXYL stearic acid. The data points for the gating arginines are shown in red boxes. To the right of each graph, the strength of the paramagnetic relaxation is mapped onto the average paddle NMR solution structure, with residues most strongly affected in red and those least affected in blue. Unassigned residues are shown in green.

The third gating charge in the NMR average structure is close to serine 1609 on the S3b helix with a distance between the serine oxygen and one of the terminal arginine nitrogens of 3.5 Å (Figure 6), suggesting that the two residues interact with each other. In the activated state, it is thought that an acidic residue on the S2 helix interacts with the third gating charge to stabilize the helix^13^. However, our observation of the close proximity of S1609 to R3 suggests that this interaction may also contribute to the stabilization of R3 in the activated state. Interestingly, the only currently known point mutation on the NaV1.5 DIV S3b helix to cause a disease phenotype is this same residue, serine 1609. This mutation, S1609W, causes long QT syndrome type 3 (LQT3)^44^, but to the best of our knowledge the functional effect of this mutation on NaV1.5 has not been studied in detail. Based on our NMR solution structure, a tryptophan replacing this serine would disrupt the helix-helix interaction since tryptophan is much bulkier than serine. Another possibility is that a cation-pi interaction between the tryptophan and R3 might help stabilize the VSD in the activated conformation, thus explaining the gain-of-function phenotype observed in LQT3.

### Loop residues

The loop connecting the S3 and S4 helices is short and rich in aromatic residues, consisting of a tyrosine, two phenylalanines, a serine and a proline (YFFSP). The tyrosine is actually the last helical residue of the S3b helix but can be considered part of the loop due to involvement of its side chain in an aromatic ‘cap” that marks the end of S3 (Figure 6). The rings of the first two aromatic residues in this “cap” (Y-F) are facing away from S4 and are conserved throughout all VGSC subtypes. The third residue in the loop, phenylalanine 1617 points to the opposite side from the previous two aromatic residues. This residue is unique to Na_V_1.5, while in all other subtypes it is replaced by a valine.

Interestingly, deletion of the third aromatic residue (F1617del) has been reported to be associated with LQT3. Chen *et al*. discovered that this deletion alters the voltage dependence of fast inactivation through a reduction of gating charges^45^. They postulated that the shortening of the loop caused by the mutation would reduce the flexibility of the S4 helix and make it difficult for the voltage sensor to return to a fully closed, resting position during hyperpolarization and to a fully extended state during depolarization. Given the very short nature of this loop (even shorter than the 7-10 residues that Chen and co-workers expected), such an effect is indeed plausible. A possible additional effect is that loss of one of the aromatic cap residues that likely interact with the lipid head-groups could also weaken the positioning of the end of S3 with respect to the membrane, which might further disrupt the correct conformational shifts in S4 upon membrane potential changes. Interestingly, F1617 and the following residue (S1618) are highly susceptible to paramagnetic relaxation by hydrophobic relaxation agents (see below, and Figure 7), indicating their strong interaction with the membrane lipids. Consistent with this strong lipid interaction, the two residues are barely affected by the water-soluble relaxation agent Mn^2+^ even though they are part of a potentially solvent-exposed loop.

### Interaction with micelle

To determine the residues that are inserted into the hydrophobic core of the micelle and the ones that reside at the micelle-solvent interface, we measured paramagnetic relaxation caused by water-soluble Mn^2+^ ions as well as two hydrophobic spin-labeled reagents, 16-doxylstearate and 5-doxylstearate (Figure 7). Residues located deeper in the hydrophobic interior of the micelle are expected to be affected more strongly by 16-doxylstearate, while those closer to the surface by 5-doxylstearate. Overall, the strongest relaxation effects were observed in samples with 5-doxylstearate, which reduced the intensity of most residues at a 1:2 peptide to spin label molar ratio. The strongest effect of 5-doxylstearate was on residues V1599, I1600, S1602, G1605 and T1606 on the S3b helix and on R1626, R1629, I1630 and G1631 on the S4a helix). The most protected residues were those at or near the loop region, from V1607 on S3b to F1616 in the loop and from L1621 to I1625 on the S4 helix. As indicated above, the exceptions are residues F1617 and S1618, which are strongly affected by the spin label in spite of their position in the loop. A similar pattern is seen with 16-doxylstearate, however the effect of 16-doxylstearate is generally weaker than 5-doxylstearate. This shows that the peptide is inserted into the micelle but generally resides in the more radial regions of the hydrophobic core. The unusual behavior of the loop residues F1617 and S1618 may be due to strong lipid interactions as stated above, and possibly also due to specific interactions of these two side chains with the DOXYL group itself. The results of the paramagnetic relaxation with Mn^2+^ show that the most solvent-accessible region of the paddle domain is found towards the end of the S3 helix and specifically around residue D1610. It is possible that the relaxation effects around that residue could also be reinforced by specific interactions between the manganese ion and the aspartate side chain. However, it is interesting to note that this region of the S3 helix was shown to interact with the scorpion toxin Lqh3 by cryo-EM^46^, and therefore would be a region that is expected to be more solvent accessible to allow for the toxin binding to occur. The loop itself is not very strongly affected, presumably due to its highly aromatic character and strong interaction with the lipid headgroups. The other region that is strongly affected by the manganese are the last few residues at the C-terminal end of the fragment. This region was found to be mostly unstructured and highly flexible in our truncated construct. This dynamic behavior probably enables transient conformations where the terminal residues leave the micelle core and thus become susceptible to strong paramagnetic relaxation in the presence of manganese ions.

### Backbone Dynamics

NMR ^15^N relaxation experiments were conducted to explore the backbone dynamics of the isolated paddle motif in DPC micelles. *R*_1_ and *R*_2_ relaxation rates as well as heteronuclear ^1^H-^15^N NOEs were measured and S^2^ order parameters derived using the Lipari-Szabo model-free approach^32^. Based on these measurements, it can be seen that the paddle is quite rigid overall with S^2^ values above 0.9 for almost half of the peptide. Based on the order parameters, the most rigid region of the peptide is the first half of the S3b helix, with increased dynamics in the second half of S3b, the loop region and in the S4 helix (Figure 8). The most flexible part of the peptide is found to consist of the last three residues on the S4a helix, which is most likely due to the truncation and can further explain why the C-terminal region of the peptide did not converge in the structure calculations. However, it is interesting to note that the N-terminal end of the peptide, which is also truncated, does not experience the same level of dynamics, which could indicate that the level of conformational heterogeneity and dynamics at the C-terminal end of the fragment is inherent to the amino acid sequence itself. This would be consistent with the extensive conformational rearrangements that the S4 helix undergoes during voltage gating. The generally lower order parameters in the end of S3b, the loop region and S4 also demonstrate that these regions of the sequence have an inherent flexibility that allows for the structural rearrangements that the VSD undergoes during voltage gating.

**Figure 8:**
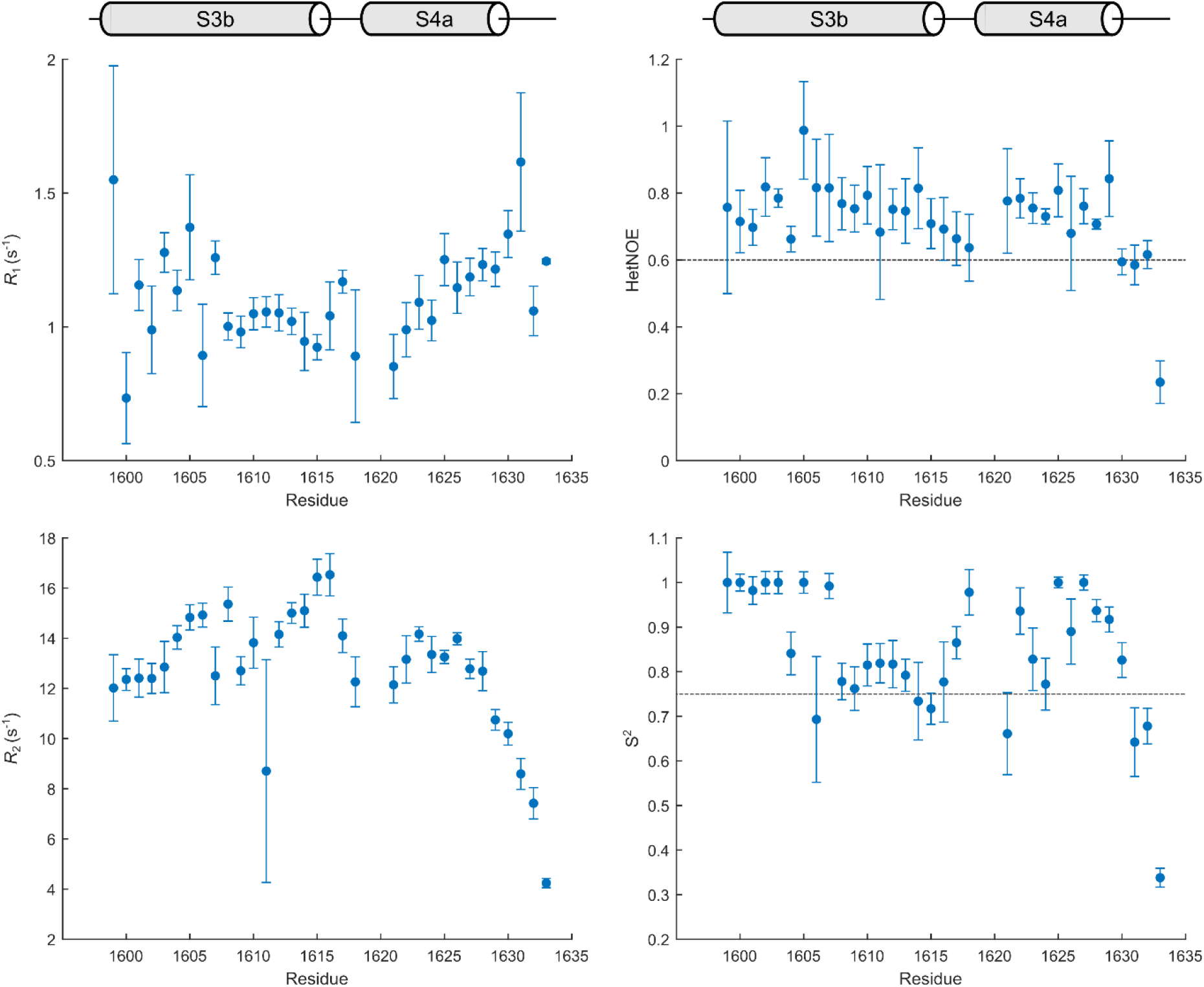
Overview of *R*_1_, *R*_2_, ^1^H-^15^N heteronuclear NOE and S^2^ relaxation parameters of NaV1.5 DIV S3b-S4 paddle motif in DPC micelles. Backbone relaxation rates were obtained on an 800 MHz spectrometer at 37 °C, and the S2 order parameters calculated through Lipari-Szabo model free analysis. Residues with HetNOE <0.6 or S^2^ <0.75 are subject to increased motion in the ps-ns time scale.

In addition to the overall trends described above, there are a few specific residues that stand out due to their increased dynamic behavior. T1606, for which we obtained the lowest order parameter in the S3 helix, is located at the position where helix S3 forms a kink. In the structure of an intermediate state of Na_V_1.5 induced through scorpion toxin binding, the kink in the helix at that position increases, facilitating the downward translocation of the S4 gating charges by two positions^46^. The increased dynamics observed for this threonine therefore appear to be essential in allowing the adjustment in the degree of helix kinking to happen. The residues lysine 1614 and tyrosine 1615 at the end of the S3b helix as well as leucine 1621 at the beginning of S4a also display S^2^ values below 0.75. These residues bracket the S3-S4 loop and therefore their increased dynamics points to a role as hinge points during the gating transitions. It is also interesting to note that backbone relaxation parameters could not be obtained for the two residues preceding L1621, proline 1619 and threonine 1620. While this is due to the absence of an amide proton for the proline residue, the fact that the amide ^1^H-^15^N correlation signal for T1620 could not be observed may indicate a chemical exchange process happening at this residue at an intermediate time scale, further supporting the notion that this region of the structure forms a flexible hinge important in the VSD gating transition. T1620 is the first residue of the S4 helix and interestingly is the site of multiple mutations that lead to both gain and loss of function phenotypes. One of these mutations is T1620K, which causes both cardiac conduction disease and LQT syndrome^47^. Since T1620 is the first residue on the S4 helix and found about one turn of the helix before R1, mutation to lysine would simply add an extra gating charge that can explain the gain of function and loss of function characteristics of this mutant at different membrane potentials^47^. More interestingly in the context of our study, the T1620M mutation was found to cause Brugada syndrome^48^. When this mutant channel was expressed in mammalian cells, it caused a slower recovery from inactivation^49^. Given the apparent conformational flexibility of T1620 that we found in our study, it is possible that a mutation to the larger and less polar methionine side chain at this position may inhibit the conformational transitions involved in the recovery from the inactivated state.

### Conclusion

We developed a method to produce and reconstitute the Na_V_1.5 DIV S3b-S4a paddle motif in DPC micelles in sufficient amount and purity for NMR study while avoiding the use of highly toxic cyanogen bromide in the protocol. The resulting NMR solution structure resulted in a well-defined helix-loop-helix motif with only the last helical turn in the S4a segment being unstructured, likely due to the truncation. The average NMR structure aligns well with the corresponding segments of the rat and human Na_V_1.5 cryo-EM structures and provides some additional structural detail given the lower resolution of the cryo-EM structures in the VSD region. In particular, our NMR structure provides plausible structural explanations for three disease-causing mutations, S1609W, F1617del and T1620M. We then studied the interaction of the paddle motif with micelle detergent using paramagnetic agents and determined that the C-terminal end of the S3 helix is the most solvent-exposed part of the structure, whereas the rest of S3 and S4 (with the exception of the unstructured terminus) are mostly embedded in the micelle. The dynamics of the S3b-S4a paddle motif were also studied using NMR relaxation, revealing increased motions in residues expected to be involved in the structural rearrangements that happen during voltage sensing.

Based on these results, our system should be suitable for future ligand binding studies in particular with animal toxins that are known to interact with this region of the channel. For example, Na_V_1.5 DIV is targeted by several sea anemone toxins, and using NMR spectroscopy to study their interaction with the isolated paddle motif should help elucidate structural details of the toxin-channel complex. The same approach could also be used for studying the S3b-S4a motifs of the other VSDs of the cardiac as well as other sodium channels. While the latest cryo-EM structures have provided a wealth of structural insight into sodium channel structure and interactions, additional methods for higher resolution and more targeted elucidation of highly flexible regions like the one we have described here will be necessary to complement these studies.

## ACCESSION CODES

Human Na_V_1.5: Q14524. The NMR structure of Na_V_1.5 DIV S3b-S4a in DPC micelle was deposited in the PDB (**7L83**), and NMR assignments as well as relaxation values are available through BMRB (**30834**).

## Author Contributions

AKH and SFP planned the research. AKH and MHB carried out the experiments. BA and JZ contributed to peptide purification and NMR experiment setup, respectively. AKH and SFP wrote the manuscript. All authors approved the final manuscript.

## ACKNOWLEDGEMENTS

The authors acknowledge Drs. Michael Goger and Shibani Bhattacharya for assistance with NMR data collection, Dr. Leah Cohen for assistance with HPLC purification and Dr. Fred Naider for helpful discussions and instrument use.

## ASSOCIATED CONTENT

**Supporting Information**. One PDF file with figures S1-S5.

