## Supplementary Figures S1-S5 for "NMR solution structure and analysis of isolated S3b-S4a motif of repeat IV of the human cardiac sodium channel"

**[2]PhD Program in Biochemistry, The Graduate Center, CUNY, 365 Fifth Avenue, New York, NY 10016, USA**

**[3]PhD Program in Chemistry, The Graduate Center, CUNY, 365 Fifth Avenue, New York, NY 10016, USA**

### Supplementary figures

```
a gat ctc gac gac gac gac atg gtg gtt gtg atc ctg agc atc gtg ggc acc gtg ctg
   D   L   D   D   D   D   M   V   V   V   I   L   S   I   V   G   T   V   L
                                          1600

  agc gac atc atc cag aag tac ttc ttc agc ccg acc ctg ttc cgc gtt atc cgc tta
   S   D   I   I   Q   K   Y   F   F   S   P   T   L   F   R   V   I   R   L
      1610                                    1620

  gcc cgc atc ggc cgt atc tga ctc gag
   A   R   I   G   R   I   Ter
          1630
```

**Figure S1**: Sequence details of the PCR fragment used for the generation of the pET32-Na$_V$1.5DIVS3bS4a expression plasmid. The DNA sequence of the PCR fragment between the BglII (in green) and the XhoI (in blue) restriction sites is shown in lowercase letters, whereas the resulting translated amino acid sequence is shown in uppercase letters. The sequence shown in orange codes for a DDDD linker in the final fusion protein which is followed by the methionine in red where the platimun complex cleavage is achieved. The resulting paddle peptide sequence is shown in bold, and the stop codon in gray. The peptide sequence numbering corresponds to the numbering in the full channel. This fragment was ligated into the pET-32b(+) plasmid (Novagen) using the indicated restriction sites, resulting in a fusion protein consisting of thioredoxin followed by a 6xHis tag, an S-Tag and the insert shown in the figure above.

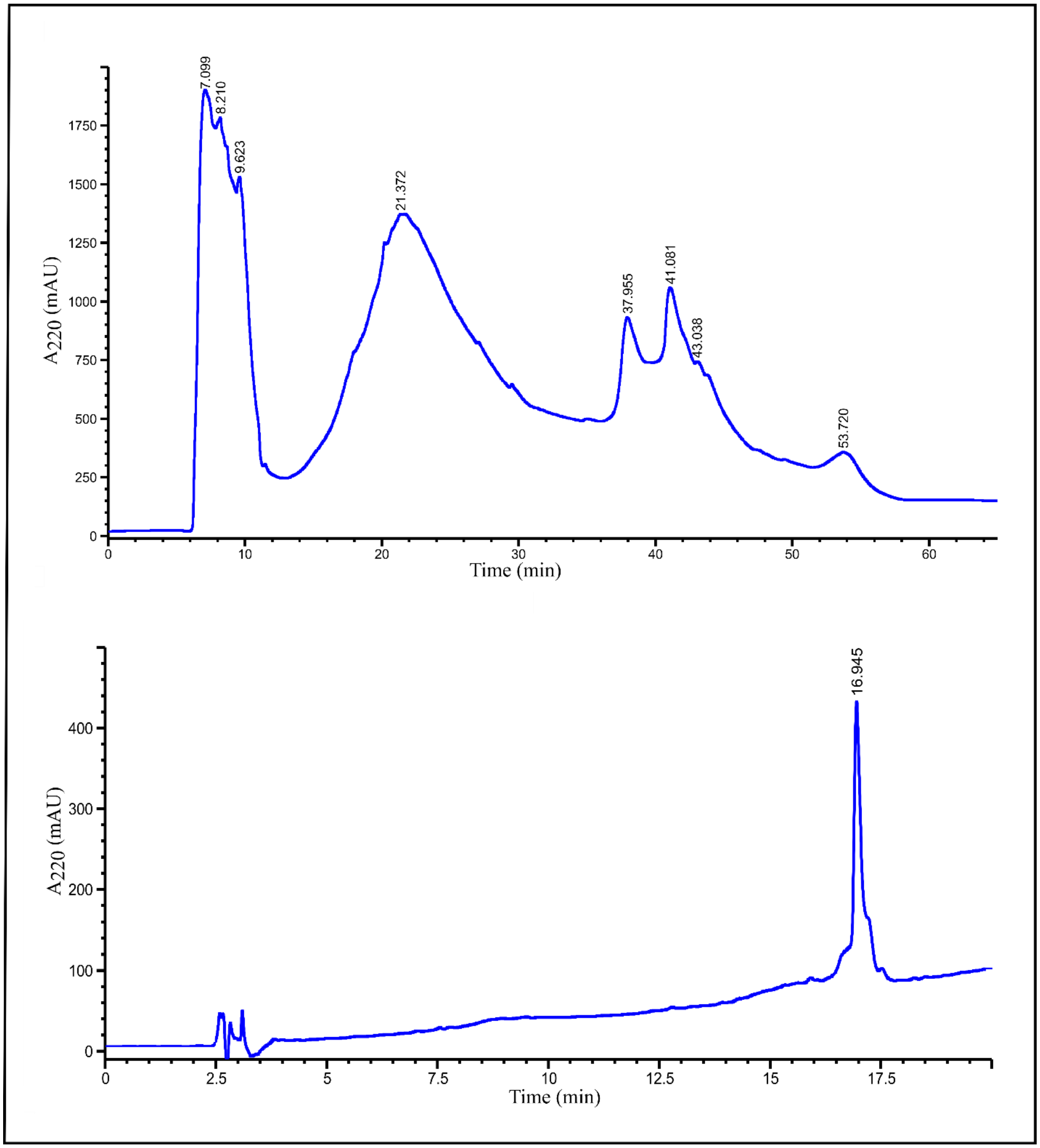


**Figure S2**: HPLC purification of the paddle motif. (top) Preparative HPLC chromatogram of the purification of paddle motif using a C3 reverse phase prep column. The paddle motif eluted at 41 minutes. Multiple fractions were collected and analyzed on MALDI-TOF. (bottom) Analytical reverse-phase HPLC chromatogram of the pure fractions as determined by mass spectrometry using a C8 column. The purity was above 85%.

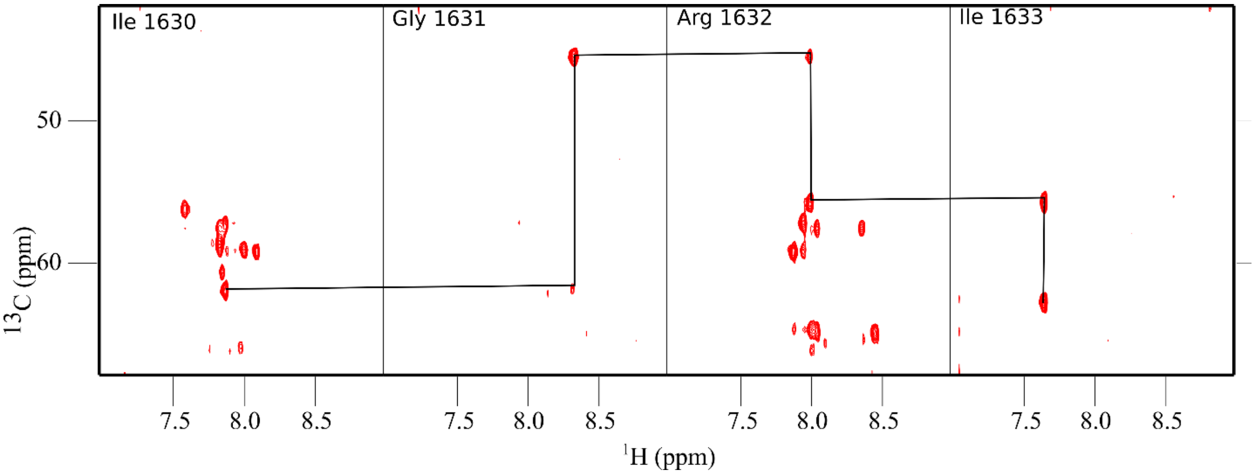


**Figure S3**: Representative strips from a 3D HNCA spectrum showing the connectivity of paddle residues 1630-1634 in Fos-Choline-10. For every strip, the $^{13}$C chemical shift of the residue's own Cα carbon is stronger than the Cα of the preceding residue, allowing the sequential assignment of the strips. Additionally, the Cα chemical shift pertaining to the preceding amino acid was also confirmed using an HNCOCA spectrum (not shown).

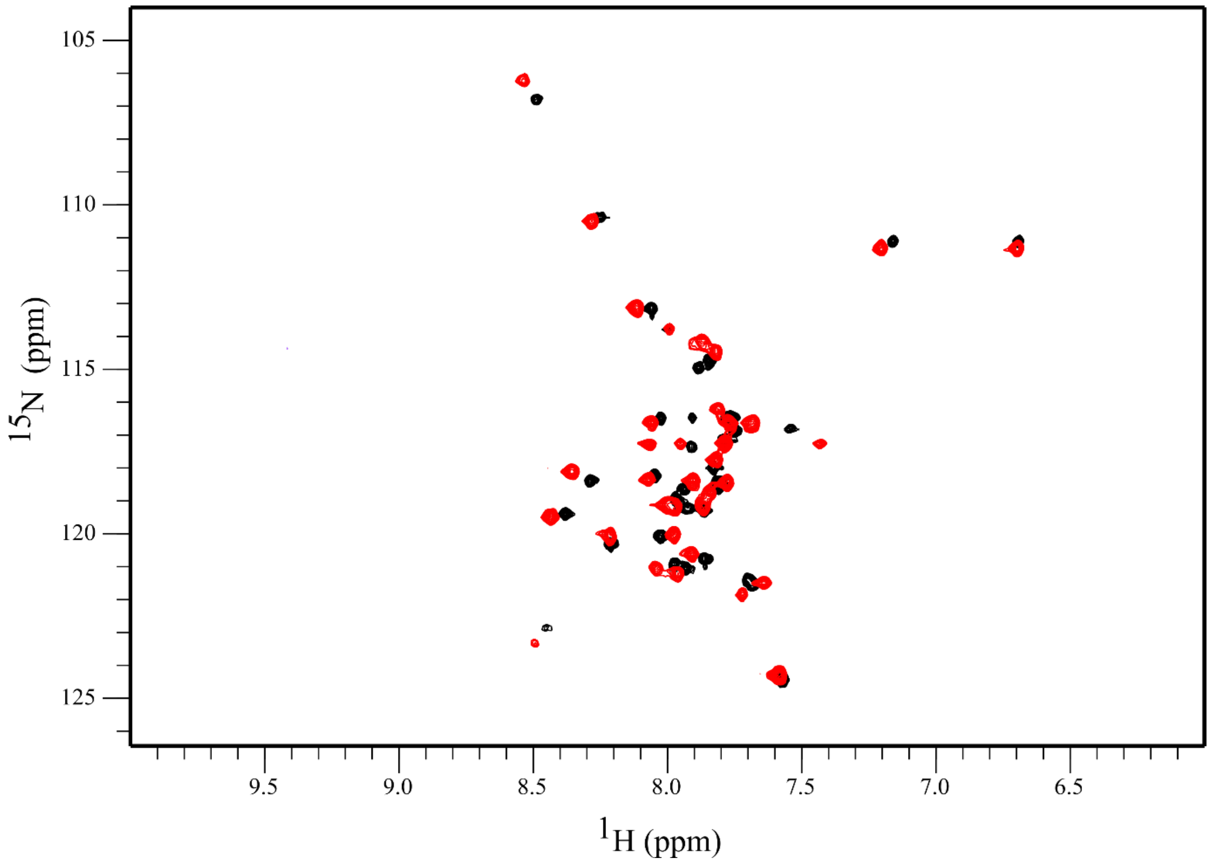


**Figure S4**: Overlay of $Na_V$1.5 DIV S3b-S4a paddle motif in DPC (red) and Fos-Choline-10 (black) showing the high level of similarity between the two spectra. This indicates that the paddle motif adopts a highly similar conformation in these two detergent systems.

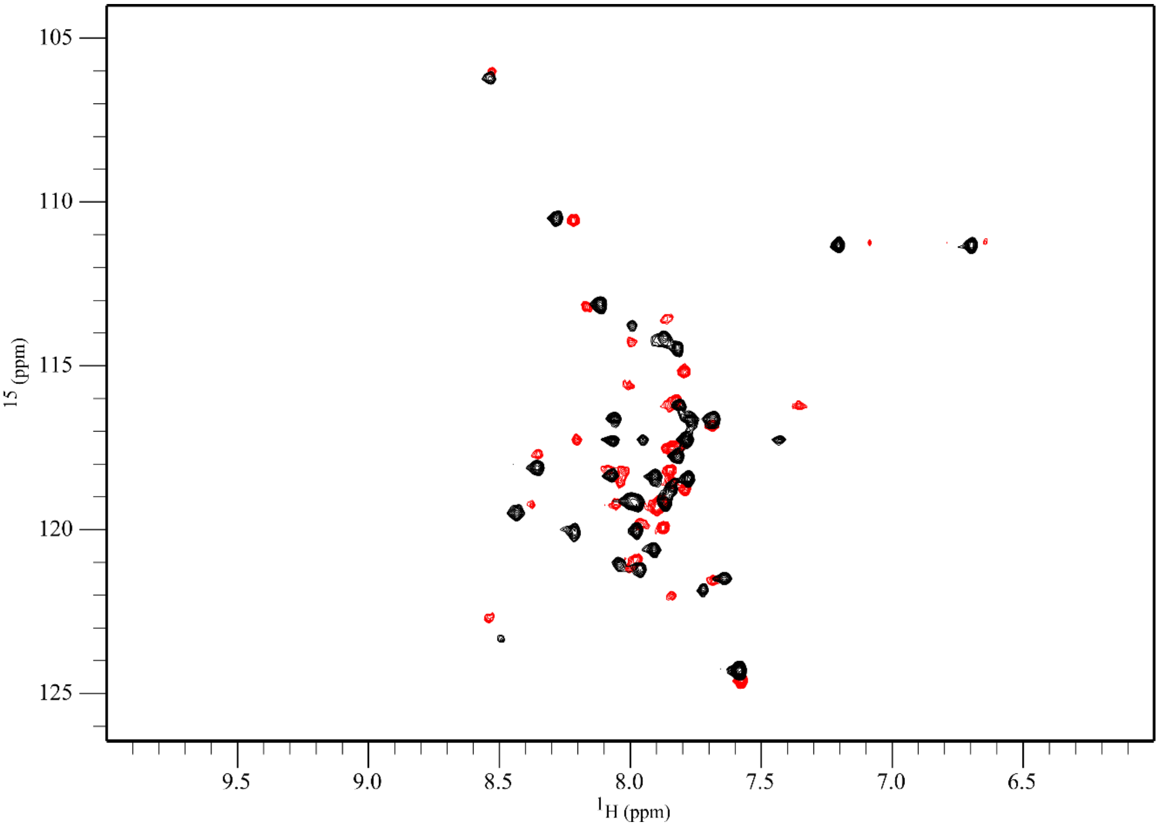


**Figure S5**:  Overlay of Na$_V$1.5 DIV S3b-S4a paddle motif in DPC (black) and DHPC/DMPC bicelles (red). The high level of similarity between the two spectra indicates that the conformation of the paddle fragment in DPC micelles closely matches that in the more membrane-like phospholipid bicelle system.
